# Draft genome of the invasive coral *Tubastraea* sp.

**DOI:** 10.1101/756999

**Authors:** Giordano Bruno Soares-Souza, Danielle Amaral, André Q. Torres, Daniela Batista, Aline Silva Romão-Dumaresq, Luciana Leomil, Marcela Uliano-Silva, Francesco Dondero, Mauro de Freitas Rebelo

**Affiliations:** Bio Bureau Biotechnology, Rio de Janeiro, RJ, Brazil; Biophysics Institute Carlos Chagas Filho, Federal University of Rio de Janeiro, RJ, Brazil; SENAI Innovation Institute for Biosynthetics, National Service for Industrial Training, Center of the Chemical and Textile Industry (SENAI CETIQT), Rio de Janeiro, RJ, Brazil; Berlin Center for Genomics in Biodiversity Research (BeGenDiv), Berlin, Germany; Università degli Studi del Piemonte Orientale (UNIPO)

## Abstract

Corals have been attracting huge attention due to the impact of climate change and ocean acidification on reef formation and resilience. Nevertheless, some coral species have been spreading very fast, replacing native species and affecting local biodiversity. Despite some focal efforts to understand the biology of these organisms, they remain understudied at the molecular level. This knowledge gap hinders the development of cost-effective strategies for management of invasive species. Here, we present the first *Tubastraea* sp. genome in one of the most comprehensive biological studies of a coral, that includes morphology, flow cytometry, karyotyping, transcriptomics, genomics, and phylogeny. The *Tubastraea* sp. genome is organized in 23 chromosome pairs and has 1.4 Gb making it the largest coral and Cnidaria genome sequenced to date. The hybrid assembly using short and long-reads has a N50 of 180,044 pb, 12,320 contigs and high completeness estimated as 91.6% of BUSCO complete genes. We inferred that almost half of the genome consists of repetitive elements, mostly interspersed repeats. Gene content was estimated as about 94,000, a high number that warrants deeper scrutiny. The *Tubastraea* sp. genome is a fundamental study which promises to provide insights not only about the genetic basis for the extreme invasiveness of this particular coral species, but to understand the adaptation flaws of some reef corals in the face of anthropic-induced environmental disturbances. We expect the data generated in this study will foster the development of efficient technologies for the management of corals species, whether invasive or threatened.

## 1. SEQUENCING THE GENOME IS A LANDMARK IN THE HISTORY OF A SPECIES

Corals are among the planet’s most stunningly beautiful organisms and they exist in an amazing variety of shapes, sizes and colors. Corals reproduce sexually or asexually [1], live alone or in colonies, with or without symbionts [2]. Corals form blooming reefs in ecosystems that range from low productivity shallow hot waters to nutrient-rich banks in the coldest depths of the ocean floor [3]. As enchanting as they are, very little is known about these amazing creatures at the molecular level. To date only seven coral species have had their genomes deposited in the NCBI Genome database (see Table 1 and Table 2).

**Table 1.**
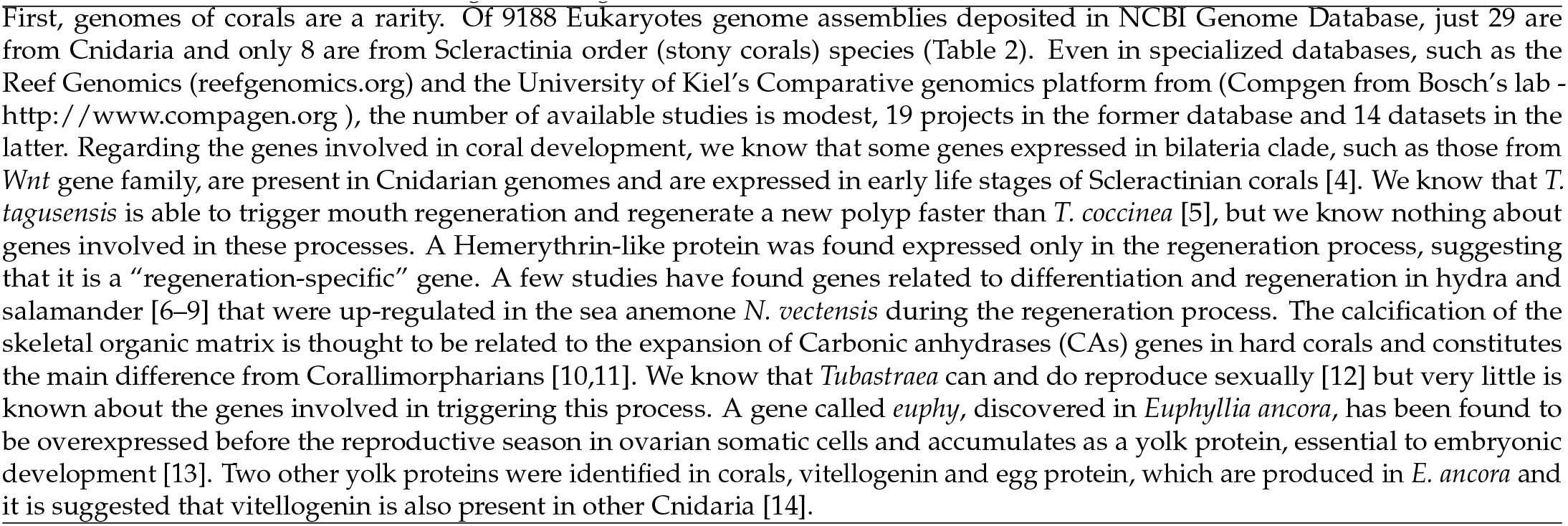
What do we know about coral genes thought to confer invasiveness?

**Table 2.**
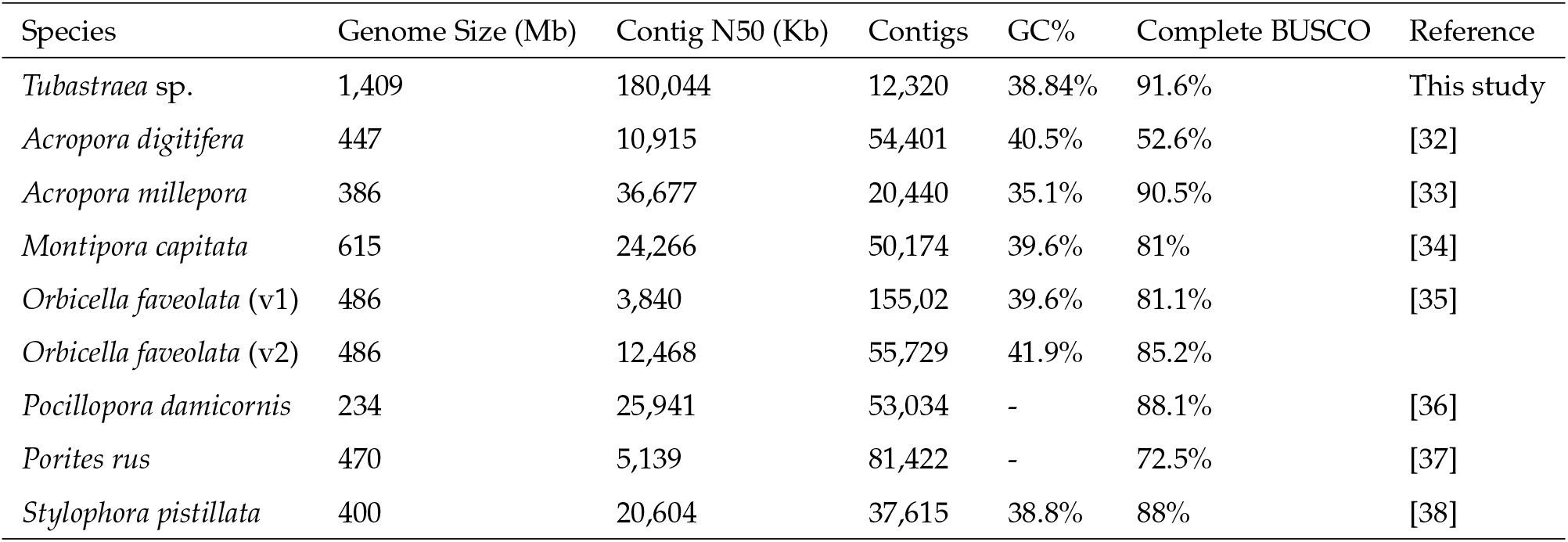
Statistics overview from our haploid draft assembly compared to other coral genomes available in NCBI’s Genbank. The *Tubastraea* sp. draft genome has the largest genome size and the highest percentage of BUSCO retrieval, making it one of the best coral genomes available so far.

This lack of knowledge hinders the development of smart strategies for both coral conservation and management of invasive coral species. The impact of climate change on coral reefs has dominated the research agenda while invasive coral species remain understudied, even in light of their significant role in the loss of biodiversity in rocky shores. *Tubastraea* sp. is a fast growth [15] azooxanthellate coral, with no substrate specificity [16], ability to produce planulae both sexually and asexually [17,18] as well as the ability to fully regenerate from undifferentiated coral tissue [5]. Since the first reports in the late 1930s in Puerto Rico and Curacao [19,20], *Tubastraea* corals (Cnidaria: Dendrophylliidae) – coral species native to the Indo-Pacific Ocean [21] – have spread rapidly throughout the Western Atlantic Ocean. Invasive *Tubastraea* corals are found in the Caribbean Sea, the Gulf of Mexico and have been detected discontinuously along 3850 km of the coast of Brazil (from 2°30’S to 26°30’S) [20,22–28] occupying up to 95% of the available substrate in some regions [12]. Recently *Tubastraea* corals were found around Eastern Atlantic islands including the Canary Islands [29,30] and Cape Verde [31]. Without innovation in control methods, dispersion is expected to continue, as desiccation in drydocks and physical removal cannot be applied in a timely and cost-effective manner, or risks inadvertently contributing to further dispersion. We know that gene-environment interactions often result in alterations in gene expression. Thus, characterization of the *Tubastraea* sp. genome should help to elucidate the molecular mechanisms of tolerance, resistance, susceptibility and homeostasis that could lead to better conservation strategies for corals as well as specific strategies to control this invasive species. The mitochondrial genome assembly could also help clarify gaps and inconsistencies in our current understanding of the evolutionary history of the genus. Here, we present the draft genome of *Tubastraea* sp., assembled using short and long reads, aggregated with RNA-seq data, flow cytometry and karyotype information and morphological characterization of the colony. In comparison to other corals it is the largest genome sequenced to date, and one of the most comprehensive efforts to elucidate genomic organization in a coral species.

## 2. WE SEQUENCED A *Tubastraea* sp. COLONY FROM SOUTHEAST BRAZIL

After collection in Angra dos Reis (23°3.229’S; 44°19.058’W - SISBIO collection authorization number 68262), the *Tubastraea* sp. colony was maintained in our laboratory. All nucleic acid extractions were performed in clone polyps. Colony morphology was analyzed and the skeleton was preserved in order to be deposited in the zoological collection of the National Museum of Brazil. The phaceloid colony measured 9.1 mm in diameter. Yellow polyps were connected by yellow coenosarc and corallites which project up to 40 mm above the coenosteum. The mean diameter of calices was 10 mm(Figure 1). In light of recent concerns raised by Bastos et al. (personal communication) regarding the misidentification of this morphotype as *T. tagusensis*, we decided to identify it only at genus level pending further analysis using both traditional and molecular techniques.

**Fig. 1.**
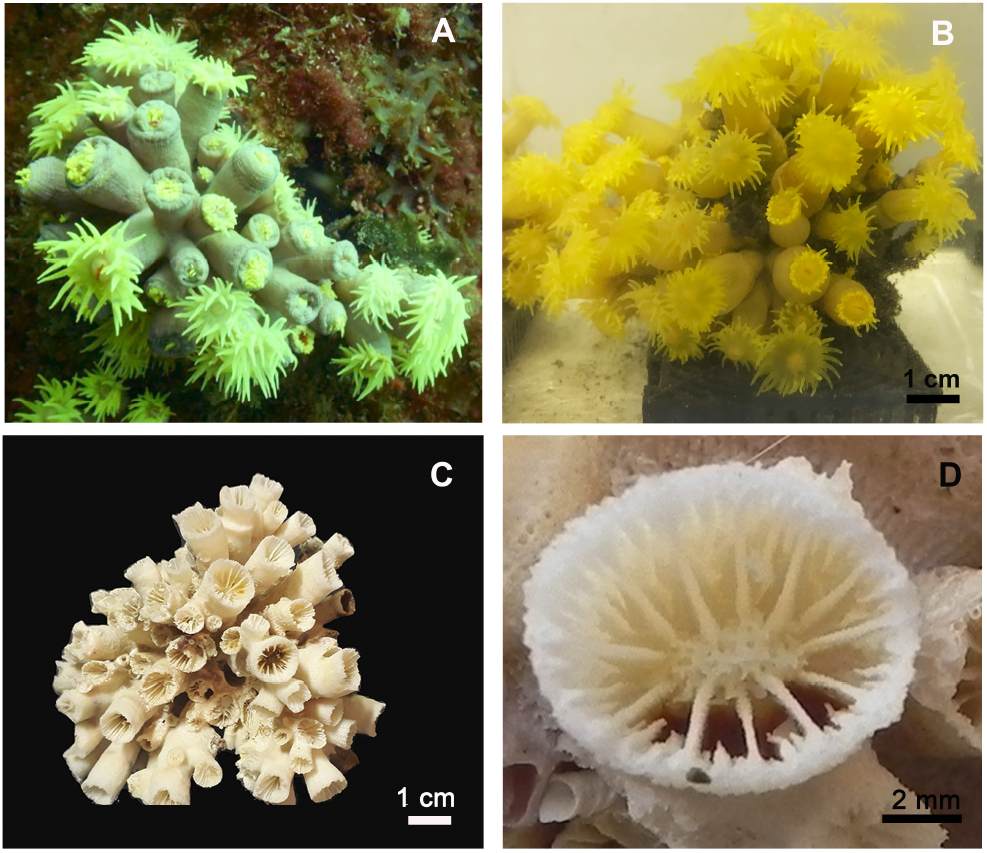
Specimen sampled for DNA extraction and sequencing. (A) Colony on substrate before removal; (B) In the laboratory, the acclimated colony in aquarium; (C) Skeleton of *Tubastraea* sp.; (D) Detail of septa arrangement, with S1 and S2 fused with columella.

## 3. *Tubastraea sp*. HAS A LARGE GENOME (1.4 GB) DIVIDED INTO 23 CHROMOSOMES PAIRS

Using flow cytometry of the nuclear suspension of a 1 mm clone polyp stained with propidium iodide, with the common pea (*Pisum sativum*) as an internal standard (See item A.2 in MM), the haploid genome size of *Tubastraea* sp. was estimated at 1.3 gigabases. The representative karyogram was assembled from one metaphase, using genetic material from planulae of the same colony treated with 0.1% colchicine, and shows a diploid karyotype of 46 chromosomes (Figure 2 and item A.3 in MM). Further analysis is necessary to identify whether there are sex chromosomes. To our knowledge, this is the first Dendrophylliidae genomic study to present evidence about the number of chromosomes and ploidy.

**Fig. 2.**
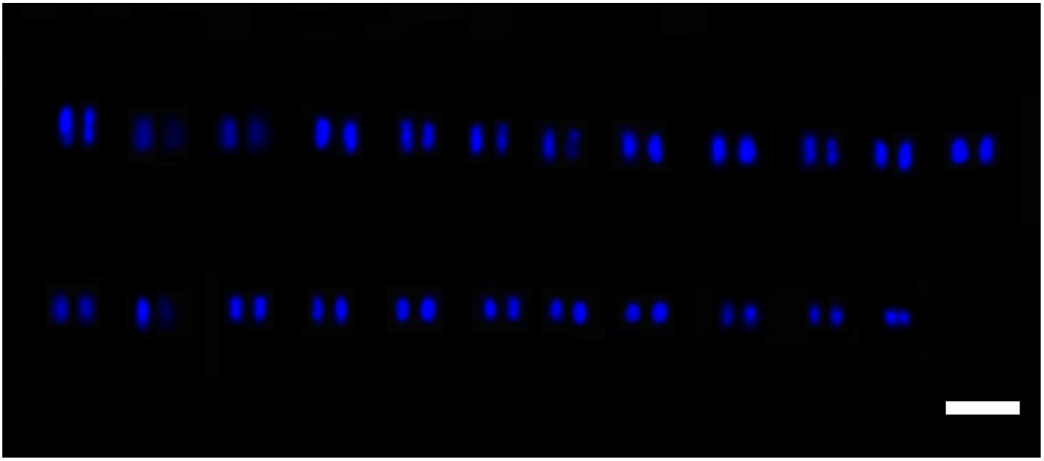
Representative karyogram of *Tubastraea* sp. from Angra dos Reis, RJ, Brazil. Bar: 5 μm.

DNA was extracted using protocols which were modified to increase the integrity (see item A.4 in MM). DNA extraction using the *DNaesy* Blood Tissue kit yielded a mean length of 10 kb with a DNA Integrity Number (DIN) of 6.4, while extraction using CTAB buffer yielded a mean length of 23 kb, both free of proteins and other contaminants. A total of 383 Gb of paired-end DNA data were obtained using an Illumina HiSeq X. The PacBio library, made skipping the shearing step, generated an average library size of 15 kb, that was subsequently sequenced on a PacBio Sequel platform and generated 54 Gb of sub-reads. RNA was extracted using the TRIZOL method according to manufacturer’s instructions. The RNA integrity number (RIN) was estimated as 6.6 (see item A.4 in MM). NEBNext mRNA libraries were built using magnetic isolation and 30 million 150 bp paired-end reads were sequenced using an Illumina HiSeq X.

## 4. AND WE RECOVERED 91.6% OF BUSCO COMPLETE GENES IN THE GENOME

We sequenced 2.5 billion of DNA paired-end short-reads comprising 383 billion bases and 5 million long-reads comprising 54 billion bases. Short-reads were quality trimmed and filtered retaining about 85% of both reads and bases. Using MaSuRCA assembler and Purge Haplotigs software to filter allelic contigs, we estimated the haploid genome size of Tubastraea sp. to be 1.4 Gb, corroborating the size estimated by flow cytometry, N50 of 180,000 kb in 12,000 contigs and completeness, as measured by BUSCO, of 91.6%, making it one of the best coral genomes made available to date.

## 5. WE RECOVERED 98.1% OF BUSCO COMPLETE GENES IN THE TRANSCRIPTOME

The stranded paired-end RNA sequencing raw data consisted of about 60 million reads and was quality trimmed and filtered yielding about 7 billion bases. Trinity assembled a *de novo* transcriptome with a little more than 200,000 genes and 300,000 transcripts with an average length of 800 bases and N50 of 1,400 kb. Of these reads, 95% mapped to the transcriptome and 75% mapped to the genome. We recovered 98% complete genes in a search for orthologous genes using the BUSCO metazoa database and we retrieved more than 5,000 (about 30%) near-complete protein-coding genes with reads mapping to more than 80% of the estimated protein length. The final set consisting of 158,075 transcripts included only sequences with coding and homology evidence.

Our draft assembly of *Tubastraea* sp. ranks as the largest Cnidarian genome published to date, three times longer (1.4 Gb vs 441±101) than published scleractinian genomes. Our hybrid assembly dramatically improved both contiguity and gene recovery, with the N50 increasing from about 6,000 bp to more than 180,000 bp, and BUSCO orthologous retrieval improving from about 50% (only short-reads) to more than 90%. We missed less than 8% of metazoan BUSCO genes in the genome and just 1% in the transcriptome. The scaffolds are available to download upon request at http://genomes.biobureau.com.br/. Repetitive intrinsic elements, inferred using ab-initio and homology-based approaches with RepeatModeler and RepeatMasker, constituted about 50% of bases, more than in Cnidaria, other invertebrates and birds, albeit similar to the percentages observed in reptiles and mammals. Genome annotation using Breaker2 provided an estimate of more than 100,000 genes. The high gene content, almost 5-fold the usual in Eukaryotes, could be attributed to reminiscent contamination, genome fragmentation, or genome duplication, and warrants deeper scrutiny. The functional annotation based on PANTHER protein families showed that the most representative pathways in Tubastraea sp. are: Wnt signaling; Integrin signalling; Gonadotropin-releasing hormone receptor pathway; Inflammation mediated by chemokine and cytokine signaling and Nicotinic acetylcholine receptor signaling. The Gene Ontology annotation showed a prevalence of protein families associated with biological regulation, cellular process localization, and metabolic process, though, unexpectedly, only a few sequences were associated with biological adhesion and cell proliferation.

## 6. THE MITOCHONDRIAL GENOME CLUSTERS WITH OTHER PHACELOID SPECIMENS

Using short and long DNA reads, we assembled a single contig containing the entire *Tubastraea* sp. mtgenome, consisting of 13 genes encoding proteins and 4 non-coding genes (Figure 3). Preliminary phylogeny of the partial sequence of the ATP8 and COI genes with several other morphotypes sampled along the Brazilian coast (data not shown) showed that *Tubastraea* sp. clusters in a monophyletic clade together with sequences of *Tubastraea tagusensis* available in NCBI databases. The lack of molecular markers from the original *T. tagusensis* samples from the Galapagos Islands leave the definitive name attribution to the species open for now.

**Fig. 3.**
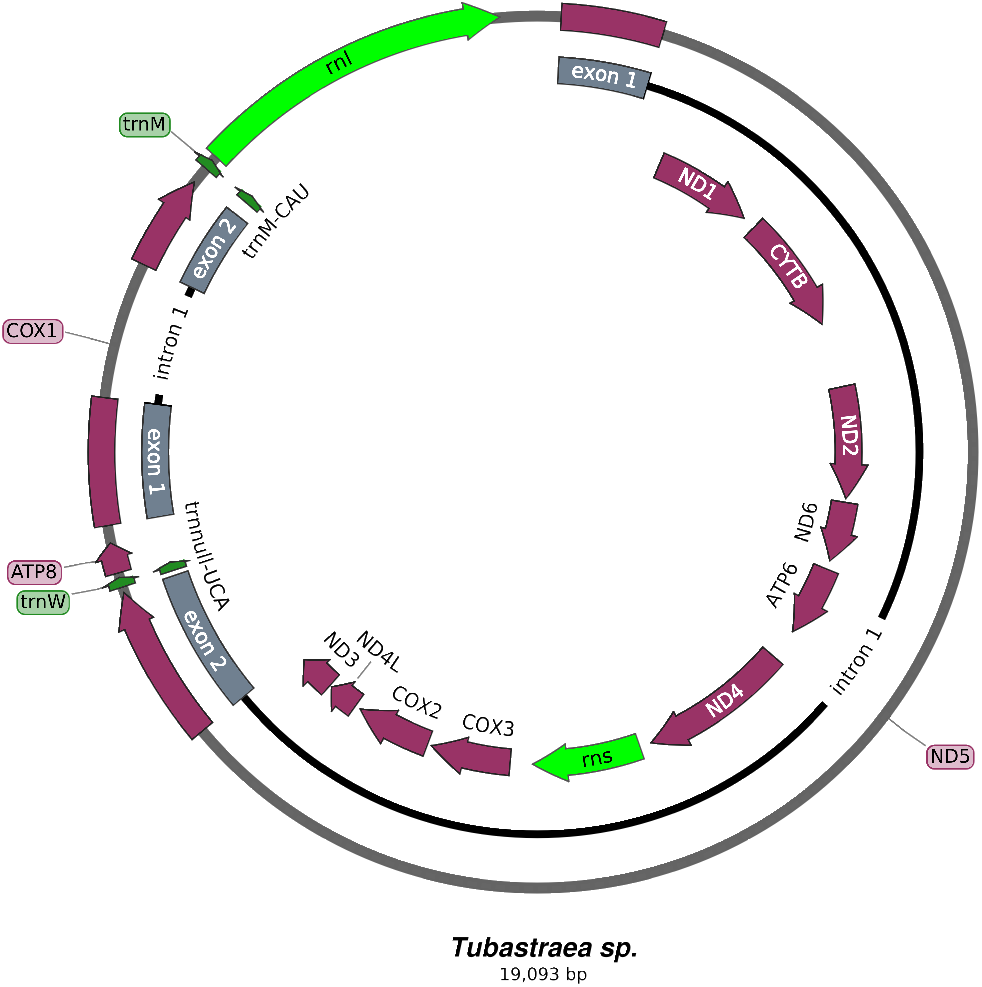
Gene map of the mitochondrial genome of *Tubastraea* sp. Genes that encode for proteins are colored in purple. The gene structures of *ND5* and **COX1** are shown parallel to the genes inside the circle. Exons are colored in grey and introns are represented by a solid black line. Genes that encode for transfer RNA and subunits of the ribosome are coloured in dark green and light green, respectively.

## 7. CONCLUSIONS

We present one of the most comprehensive studies of a scleractinian, with the morphology, cytogenetics, transcriptome, mitochondrial and draft nuclear genome of a *Tubastraea* species. It constitutes the largest genome of the Scleractinia order and Cnidaria phylum published to date. This study yielded findings which begin to fill some of the gaps in our understanding of the *Tubastraea* genus. The mitochondrial genome provides further evidence pertinent to the discussion of the species identification. Building upon the foundation of the work presented, in the next phases of our research we will improve the contiguity of the draft assembly by the use of molecular and computational scaffolding methods, elucidate the taxonomy within the *Tubastraea* genus by the use of integrative taxonomy, and generate better annotations to guide the development of biotechnological strategies to deter bioinvasion by this species of sun coral and to gain insights about resilience among native corals.

## 9. DISCLAIMER

To enhance readability, we elected to organize this article in a narrative format. The content of the conventional scientific format of Introduction, Materials Methods, Results, and Discussion is present, just not with these subtitles. A detailed materials, methods and results section is presented after the references.

## 11. ACKNOWLEDGEMENTS

Authors are indebted to Mrs Lisboa M for her support. Dr. Sousa-Marçal S for his support in karyotype assembly. Dr. Viccini LF and Dr. Monteiro E for their support in flow cytometry. Dr. Zilberberg C for her insights in DNA extraction and Guimarães M and Wajsenzon IJR for their invaluable help in experiments performace.

## A. Material, Methods and Results

### A.1. Tubastraea sp. sampling and morphological analysis

We collected a specimen of *Tubastraea* sp. by scuba diving off the rocky shores of Porcos Pequena Island in the municipality of Angra dos Reis, Rio de Janeiro state, southeast Brazil (23°3.229’S; 44°19.058’W). We carefully removed the healthy colony from the substrate and immediately transported it to the laboratory. After a period of acclimation, we transferred the colony to a 20-liter seawater-filled aquarium (pH = 8.2; T°C = 24) which were continuously aerated and subjected to a 12h:12h photoperiod. After DNA extraction, the coral skeleton was prepared for morphological analyses by placing the colony in a container filled with hypochlorite solution (NaOCl) for the removal of all soft tissues. The bleaching time was three days. Colony and coralites were photographed using a Nikon D750 digital camera and a Leica M205 FA magnifying glass. Macroscopic characteristics of colony were examined with a Leica M205 FA magnifying glass. Calipers were used to measure the diameter of the colony and calice, the intercoralite distance, and the height of the polyps. Characteristics measured in our colony were compared with published descriptions of *Tubastraea* species [21,39,40].

### A.2. Flow cytometry

The genome size of *Tubastraea* sp. was estimated by flow cytometry. For the nuclei suspension preparation, a polyp was sampled and a 1 mm piece was minced in a buffer containing 0.2 M Tris-HCl, 4 mM MgCl2.6H2O, 2 mM EDTA Na2·2H2O, 86 mM NaCl, 10 mM sodium metabisulfite, 1% PVP-10, 1% (v/v) Triton X-100, pH 7.5. *Pisum sativum* (pea) was chopped in the same buffer and used as an internal standard for genome size estimation. The nuclear suspension was stained with propidium iodide and at least 5,000 events were analyzed using CytoFLEX (Beckman Coulter Life Sciences). Histograms were analyzed using CytExpert 2.0 software (Beckman Coulter Life Sciences) (Figure 4). The 2C DNA content of *Tubastraea* sp. was 2.61 (± 0.78) picograms, equivalent to 2.5 Gb (± 0.076) and was calculated as the sample peak mean, divided by the *P. sativum* peak mean and multiplied by the amount of *P. sativum* DNA (9.09 pg, [41]). The procedure was performed in experimental quadruplicate. The estimated genome size of the *Tubastraea* sp. specimen obtained in Angra dos Reis was 1,277.5 Mb.

**Fig. 4.**
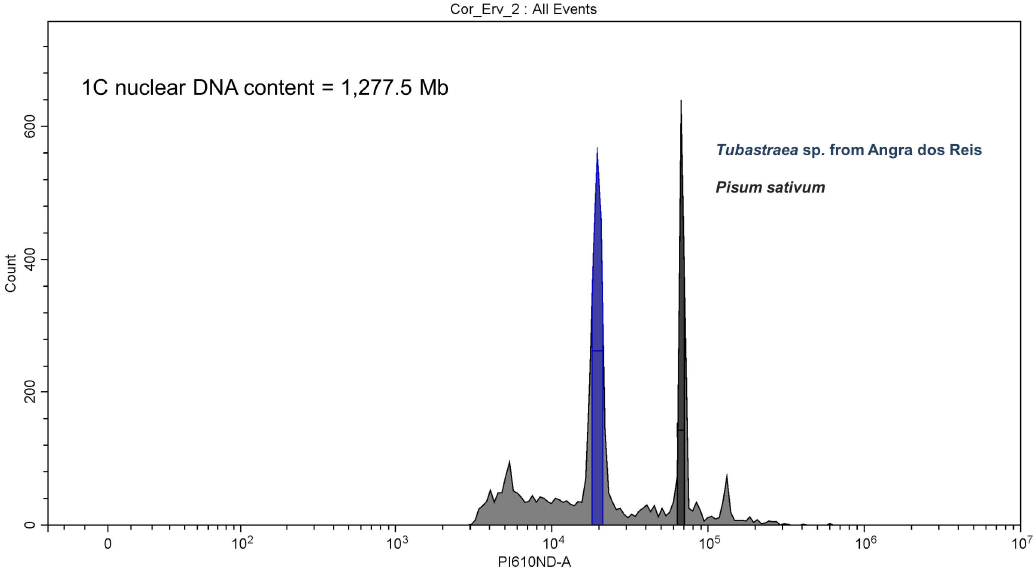
Representative histograms of DNA staining in nuclei with propidium iodide. In black, nuclear DNA fluorescence of the standard *P. sativum* (pea) and in blue fluorescence of *Tubastraea* sp. from Angra dos Reis. 2C nuclear DNA content of *Tubastraea* sp. was 2.61 (± 0.78) picograms, equivalent to 2,555 Mb (±76,81) (1 pg = 978 Mb). Considering diploidy, the genome size was estimated in approximately 1,300 Mb.

### A.3. Karyotype determination

Planulae from *Tubastraea* sp. were collected for karyotype determination. Planulae were exposed to colchicine (0.1%) for 24h, then exposed for 90 minutes to an osmotic shock with distilled water to rupture the membranes, and then fixed with Carnoy’s solution (3:1; Ethanol: acetic acid). The fixed planulae were then minced in acetic acid (60%), placed onto a heated glass slide and air-dried. Nucleic acids on the slide were stained with DAPI (4,6-diamidino-2-phenylindole) for visualization under fluorescence microscopy. Chromosomes were measured using Adobe Photoshop (Adobe Systems) and a representative karyogram was assembled with a karyotype 2n=46 chromosomes (Figure 2).

### A.4. DNA and RNA extraction, library construction and sequencing

Soft tissue from the coral specimen was collected, rinsed with distilled water and DNA extraction was performed. The DNA extraction from samples destined for Illumina sequencing was performed using *DNaesy* Blood and Tissue kit (Qiagen). The DNA extraction from samples destined for PacBio sequencing was performed according to Garcia and collaborators [42], optimized with several modifications. Briefly, soft tissue from sun coral specimens retrieved from aquaria were rinsed with distilled water and immersed in 1.0 ml of CTAB buffer [2 % (m/v) CTAB (Sigma-Aldrich), 1.4 M NaCl, 20 mM EDTA, 100 mM Tris–HCl (pH8,0)], with 10 μg of proteinase K (Invitrogen) and 2% of 2-mercaptoethanol (Sigma-Aldrich), freshly added, per 100 mg of tissue. Then, the tissue was kept in a lysis buffer for 4 days, with occasional inversion to promote tissue lysis. Tubes with tissue and buffer were then exposed to freezing with liquid nitrogen for 30 seconds and then thawed and heated to 65°C in a heat block for approximately 3 minutes. Three freeze-thaw cycles were performed. Then to remove protein and lipids one wash was performed with Phenol:Chloroform:Isoamyl Alcohol (25:24:1) (Sigma-Aldrich), and two washes with Chloroform:Isoamyl Alcohol (24:1) (Sigma-Aldrich). The supernatant was transferred to a new tube containing 1 ml of C4 solution from a Power Soil DNA Isolation kit (MO BIO Laboratories), homogenized by inversion and loaded in a spin column from the *DNeasy* Blood and Tissue kit (Qiagen, Germany). A final wash was performed with 500 μl of C5 solution. DNA elution was done with 150 μl of Tris-HCl buffer (10 mM, pH 8.5) in three sequencing centrifugation steps. RNA extraction from *Tubastraea* sp. from Angra dos Reis was done using Trizol protocol, according to the manufacturer’s instructions with a few modifications. A polyp of each species was stored in 1 ml of Trizol with 6 μl of HCl (6M) [43], overnight at 4°C. Then tissue and skeleton were homogenized with a rotor-stator centrifuge to remove debris and then we followed the Trizol manufacturer’s protocol.

Both DNA and RNA purity was assessed using a NanoDrop spectrophotometer (Thermo Fisher Scientific), the amount determined using a Qubit Fluorometer (Thermo Fisher Scientific), and DNA and RNA integrity by 1.0% agarose gel electrophoresis and the respective ScreenTape Assay using a 4200 Tapestation System (Agilent).

DNA extraction using the *DNeasy* Blood and Tissue kit yielded a DNA of about 10 kb in size with a DNA Integrity Number (DIN) of 6.4 (Figure 5) and free of proteins, according to NanoDrop spectrophotometer ratios (A260/280=1.8) which was used to construct the Illumina library.

**Fig. 5.**
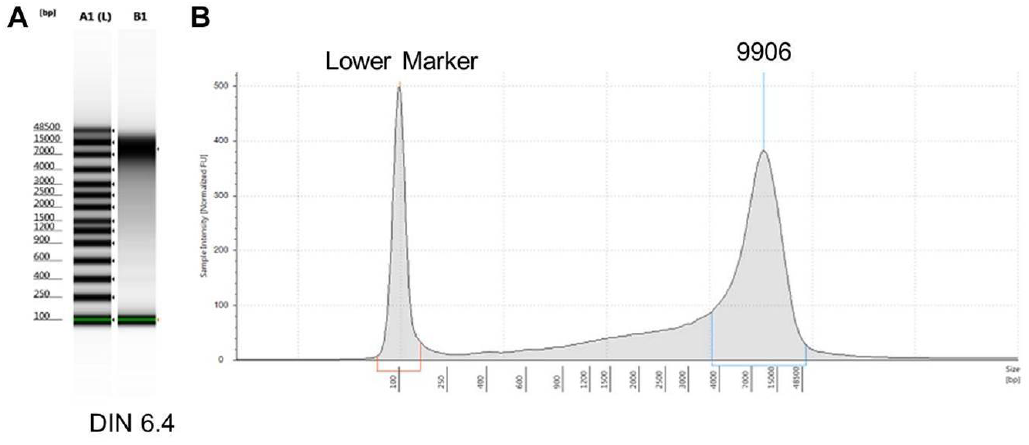
A: Electrophoresis image and DIN displayed below and (B) the electropherogram traces of genomic DNA from *Tubastraea* sp. from Angra dos Reis.

DNA extraction using CTAB buffer protocol yielded a DNA of about 23 kb in size (Figure 6) and also free of proteins and other contaminants, according to NanoDrop spectrophotometer A260/280 and A260/230 ratios, as can be seen in Table 3. The 10 samples from different elutions were pooled together before size quality control.

**Fig. 6.**
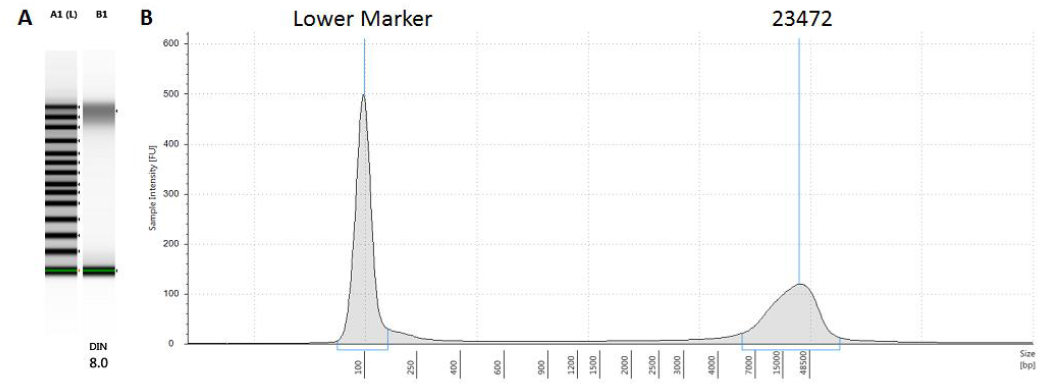
Electrophoresis image and DIN displayed below and (B) the electropherogram traces of genomic DNA from *Tubastraea* sp. from Angra dos Reis.

**Table 3.** Quantitation using Qubit BR dsDNA assay and quality evaluation using Nanodrop from different aliquots of *Tubastraea* sp. from Angra dos Reis. These samples were pooled together for SMRT bell template construction.

| DNA Aliquot | Qubit (ng/μl) | $A_{260}/_{280}$ | $A_{260}/_{230}$ |
| --- | --- | --- | --- |
| DNA *Tubastraea* sp. (1) | 286.0 | 1.89 | 2.35 |
| DNA *Tubastraea* sp. (2) | 52.7 | 1.91 | 2.24 |
| DNA *Tubastraea* sp. (3) | 27.7 | 1.82 | 2.24 |
| DNA *Tubastraea* sp. (4) | 17.4 | 1.77 | 1.51 |
| DNA *Tubastraea* sp. (5) | 455.0 | 1.86 | 2.35 |
| DNA *Tubastraea* sp. (6) | 98.6 | 1.86 | 2.19 |
| DNA *Tubastraea* sp. (7) | 34.0 | 1.83 | 2.11 |
| DNA *Tubastraea* sp. (8) | 24.5 | 1.87 | 2.06 |
| DNA *Tubastraea* sp. (9) | 700.0 | 1.86 | 2.35 |
| DNA *Tubastraea* sp. (10) | 151.0 | 1.85 | 2.34 |

RNA integrity number (RIN) was 6.6, with enough quality to proceed to Illumina library construction (Figure 7).

**Fig. 7.**
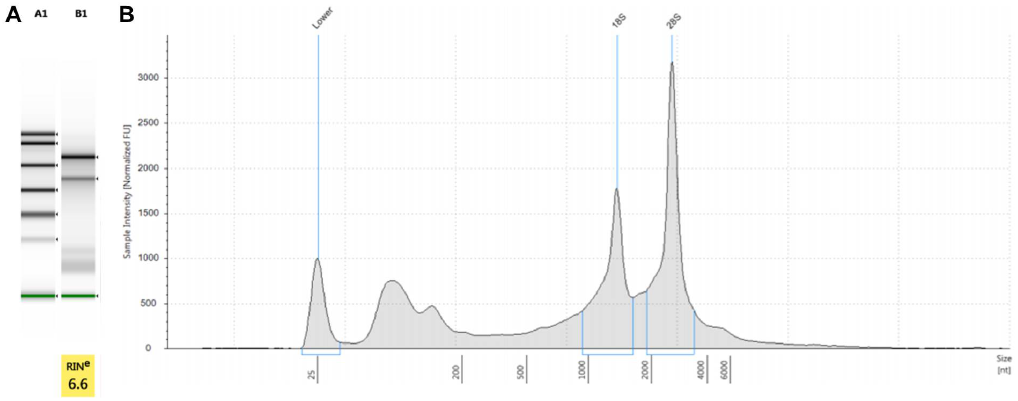
A: RNA integrity evaluation. A: Electrophoresis image and RIN displayed below and B: the electropherogram traces of the RNA from *Tubastraea* sp.

### A.5. Illumina Library construction and sequencing

The DNA library from *Tubastraea* sp. was prepared using the Kapa Hyper DNA Library Preparation Kit, with a final library size of 494 pb (Figure 8). 150 PE library was sequenced on a HiSeq X (Illumina, Inc., San Diego, US-CA). Three lanes were sequenced generating 383 Gb of raw data.

**Fig. 8.**
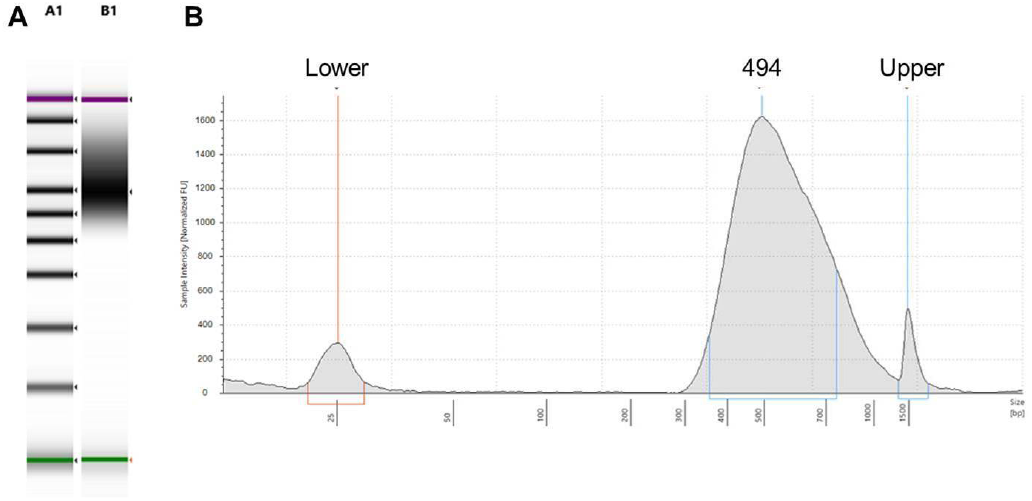
Final Library quality control. (A) Electrophoresis image and (B) the electropherogram traces of the constructed library.

### A.6. PacBio library construction and sequencing

Five micrograms of DNA from *Tubastraea* sp. were used for the PacBio library preparation. DNA were processed without further shearing as the DNA was already measured to be around 20 Kb. After DNA repair, ligate adapters, and the removal of DNA without attached SMRTbell primers, size selection was performed using Blue-Pippin (Sage science). DNA fragments in the size range >15 kb were collected and concentrated using 1xAMPure purification. The average size of the library was 15 kb (data not shown). PacBio Sequencing was carried out using the PacBio Sequel Single Molecule Real Time (SMRT) sequencing platform, Sequel in 4 SMRT cells (1M v3) generating 54 Gb.

### A.7. RNA library construction and sequencing

Library construction was performed following the manufacturer’s recommendation for the NEBNext Ultra II RNA Library Prep Kit for Illumina with NEBNext Poly(A) mRNA Magnetic Isolation Module (New England Biolabs, Ipswich, MA). Samples were pooled and sequenced on a HiSeq X sequencer at a 150 bp read length in paired-end mode, with an output of 30 million reads per sample.

### A.8. Transcriptome Quality control

The stranded paired-end RNA sequencing raw data comprised 63,515,770 reads and 9,580,881,270 bases and was first evaluated using fastQC v.0.11.8 and KAT v.2.4.1. After the visual inspection of the graphs and statistics generated, bbDUK v.38.42 was used to perform the quality trimming and filtering of reads. Illumina PE data was modulo trimmed (ftm = 5) due to the lower quality of the last base in the 151 bp reads. In the second round, reads were quality trimmed using the following parameters: i) min-length=100 (minimum length); ii) ktrim=r (trim reads matching reference k-mers); iii) k=23 (k-mer size used to find contaminants); iv) hdist=1 (Hamming distance for reference k-mers); v) tbo=t (trim adapters based on reads overlap); vi) tpe=t (trimming on both reads); vii) qtrim=rl (trim both reads’ ends based on quality score); viii) trimq=20 (quality trimming score); and ix) minavgquality=20 (minimum average read quality). Read contaminants were searched against phix and human sequences. Upon completion of these quality control procedures 26,289,606 paired-end reads and 7,347,464,831 bases were retained.

### A.9. Transcriptome Assembly

Trinity v.2.8.5 was used in the *de novo* transcriptome assembly of *Tubastraea* sp. with the following parameters: –seqType fq -max_memory 50G –CPU 16. The quality of transcriptome assembly was evaluated by different metrics such as: i) transcriptome statistics; ii) reads mappability in the transcriptome and the genome; iii) proportion of full-length reconstructed proteincoding genes from predicted transcripts; iv) proportion of recovered conserved orthologous genes. Trinity v.2.8.5 stats script was used to calculate the summary statistics of transcriptome assembly. 208,419 genes and 357,670 transcripts were inferred from RNA-seq data with a transcript contig N50 of 1,407 and transcript average length of 803.13. We estimated 94.81% of the reads were mapped to the transcriptome using bowtie2 v.2.3.5 with the following parameters (-p 16 -q –no-unal -k 20 -x) and 75% of them aligned to the genome using STAR v.2.7.2a with default parameters. To estimate the number of proteincoding genes represented by putative transcripts we trailed the following steps: i) “blasted” (blastx v.2.9.0+) the putative transcripts against the SwissProt database release 2019_07 (-evalue 1e-20 -num_threads 16 -max_target_seqs 1 -outfmt 6); ii) estimated the proportion of protein sequence targets aligned to the best match transcript (analyze_blastPlus_topHit_coverage.pl); iii) grouped multiple high scoring segment pairs (HSPs) hits to estimate sequence coverage based on multiple alignments (blast_outfmt6_group_segments.pl); iv) compute the percent coverage by length distribution. Then, we recovered 5638 (28%) of near-complete protein-coding genes (>80% mapping length). We also computed the number of predicted orthologous genes recovered from transcriptome assembly using BUSCO v3.0.2 (Benchmarking Universal Single-Copy Orthologs) with default flags (-m transcriptome -c 32 -sp fly; metazoa database). We recovered 98.1% of complete genes; 1.3% of fragmented genes and 0.6% of missing genes.

### A.10. Transcriptome Annotation

TransDecoder.LongOrfs v5.5.0 with default parameters was used to identify the most likely candidate regions in assembled transcripts. The predicted open reading frames (ORFs) were then searched using blastp v2.9.0+ (-outfmt 6 -evalue 1e-5 -num_threads 20 -outfmt 6 -evalue 1e-5 -num_threads 20 -max_target_seqs 1) against the SwissProt database release 2019_07 and using hmmscan v3.2.1 against the Pfam database v32.0 to create a homology retention filter for TransDecoder.Predict.

### A.11. Genome Quality control

The DNA paired-end short-read sequencing generated 2,537,827,094 reads and 383,211,891,194 bases of raw data. And 5,092,127 subreads and 54,104,094,838 bases were obtained from long-read sequencing. The raw data from both experiments were examined using fastQC v.0.11.8 [44] and K-mer analysis v.2.4.1 (KAT; [45]). Long-reads data were also examined using SMRT link analysis v.6.0.0.47836 and stsPlots. After visual inspection of the fastQC and KAT results, we proceeded to the quality control trimming and filtering using bbDUK v.38.42. Illumina PE data was modulo trimmed (ftm = 5) due to the lower quality of the last base in the 151 bp reads. In the second round, reads were quality trimmed using the following parameters: i) min-length=100 (minimum length); ii) ktrim=r (trim reads matching reference k-mers); iii) k=23 (k-mer size used to find contaminants); iv) hdist=1 (Hamming distance for reference k-mers); v) tbo=t (trim adapters based on reads overlap); vi) tpe=t (trimming on both reads); vii) qtrim=rl (trim both reads ends based on quality score); viii) trimq=20 (quality trimming score); and ix) minavgquality=20 (minimum average read quality). Read contaminants were searched against the phix control library (version) and human sequences. After QC rounds, we retained 2,183,079,384 reads and 323,235,024,412 bases from Illumina PE sequencing and 5,092,127 subreads and 54,104,094,838 bases from PacBio long-reads sequencing.

### A.12. Hybrid Assembly

The genome assembly of *Tubastraea sp*. was performed with MaSuRCA v.3.3.1 (Maryland Super-Read Celera Assembler; [46]) using Flye as final assembler and default parameters apart of JF_SIZE = 30,000,000,000 and PE = pe 494 48. The quality control of *Tubastraea sp*. assembly was accessed by QUAST v5.0.2 [47] (Quality Assessment Tool for Genome Assemblies), BUSCO v3.0.2 [48] Benchmarking Universal Single-Copy Orthologs) and stats.sh from BBMap v.38.42. The inflated estimated genome size (1.8 Gb) and a high percentage of duplicated gene copies on BUSCO (76.1%) suggested the presence of diploid sequences in the first draft assembly. Then, to have a haploid assembly we used the purge_haplotigs pipeline v1.0.4 [49] by removing allelic contigs (haplotigs) from the first assembly. The final statistics from our haploid draft assembly were then compared to other coral genomes retrieved from NCBI’s Genome portal (https://www.ncbi.nlm.nih.gov/genome) (Table 2).

### A.13. Genome Annotation

Following the quality control steps, we proceeded to the annotation of the curated *Tubastraea* sp. genome. Repetitive intrinsic elements were inferred using ab-initio and homology-based approaches by RepeatModeler v1.0.11 and RepeatMasker v4.0.9 (Dfam v3.0, RepBase v20170127, and custom libraries; -gccalc -noisy -xm -xsmall -gff) (Table 4). Approximately 50.50% of bases (711,957,303 bp) were masked and the proportion of repetitive elements in *Tubastraea* sp. is similar to other Cnidaria, invertebrates and birds, albeit lower than those observed in reptiles and mammals.

**Table 4.** Summary of repeat elements found in *Tubastraea* sp. genome

| Family | Number of elements | Length occupied (bp) | % of sequence |
| --- | --- | --- | --- |
| SINEs | 116,662 | 18,789,036 | 1.33 |
| LINEs | 219,798 | 69,895,505 | 4.96 |
| LTR elements | 72,939 | 25,710,204 | 1.82 |
| DNA elements | 594,954 | 171,039,953 | 12.13 |
| Unclassified | 1,990,951 | 402,675,152 | 28.56 |
| Total interspersed repeats | 20,875 | 688,109,850 | 48.80 |
| Small RNA | 15,036 | 3,999,204 | 0.28 |
| Satellites | 295,255 | 3,901,714 | 0.28 |
| Simple repeats | 34,053 | 16,180,230 | 1.15 |
| Low complexity | 34,053 | 1,781,691 | 0.13 |

The soft-masked genome was annotated using BRAKER v2.1.3 (–cores 32 –crf –gff3 –softmasking –AUGUSTUS_ab_initio –UTR=on). BRAKER encompasses different tools to train, predict and annotate gene structures in an automated fashion. In this study, we relied on hints provided by the alignment of RNA reads to the genome using STAR v2.7.2a. GeneMark-ET, Augustus ab-initio and CRF were used to generate a set of training genes for Augustus and after that, we retained 116,715 gene models (Table 4). For gene prediction, we used BRAKER v2.1.3 with the aligned the RNA-seq bam file generated by STAR v.2.7.2a as support for gene models (–softmasking –AUGUSTUS_ab_initio –crf –UTR=on –gff3). At first, hints were generated from bam file and then processed to feed GENEMARK-ET. The gene models obtained were then filtered to be used in Augustus training. The CRF predictions performed worse than HMM and were discarded in favor of the latter. BRAKER also used Augustus for ab-initio and UTR predictions to refine gene models. The final set of genes models consisted of 10.6% of the genome covered by CDS. The summary statistics for gene models predicted in the genome are shown in Table 5. The functional annotation of predicted genes was performed using InterProScan v5.36-75.0 with Pfam v32.0 and PANTHER v14.1 as databases. The predicted genes with homology evidence were filtered to retrieve metabolic pathways and Gene Ontology annotations from PANTHER website using Panther generic mapping identifiers (Figures 9 and 10).

**Table 5.** Sumarized genome annotation statistics

| Statistics | Genome annotation |
| --- | --- |
| Number of genes | 116,715 |
| Number of mRNAs | 121,515 |
| Number of exons | 579,841 |
| Number of introns | 458,326 |
| Number of CDS | 121,515 |
| Overlapping genes | 1,061 |
| Contained genes | 439 |
| Shortest gene | 95 |
| Longest gene | 194,975 |
| mean gene length | 5,606 |
| mean mRNA length | 6,027 |
| mean exon length | 257 |
| mean intron length | 1,261 |
| mean CDS length | 1,228 |
| mean mRNAs per gene | 1 |
| mean exons per mRNA | 5 |
| mean introns per mRNA | 4 |

**Fig. 9.**
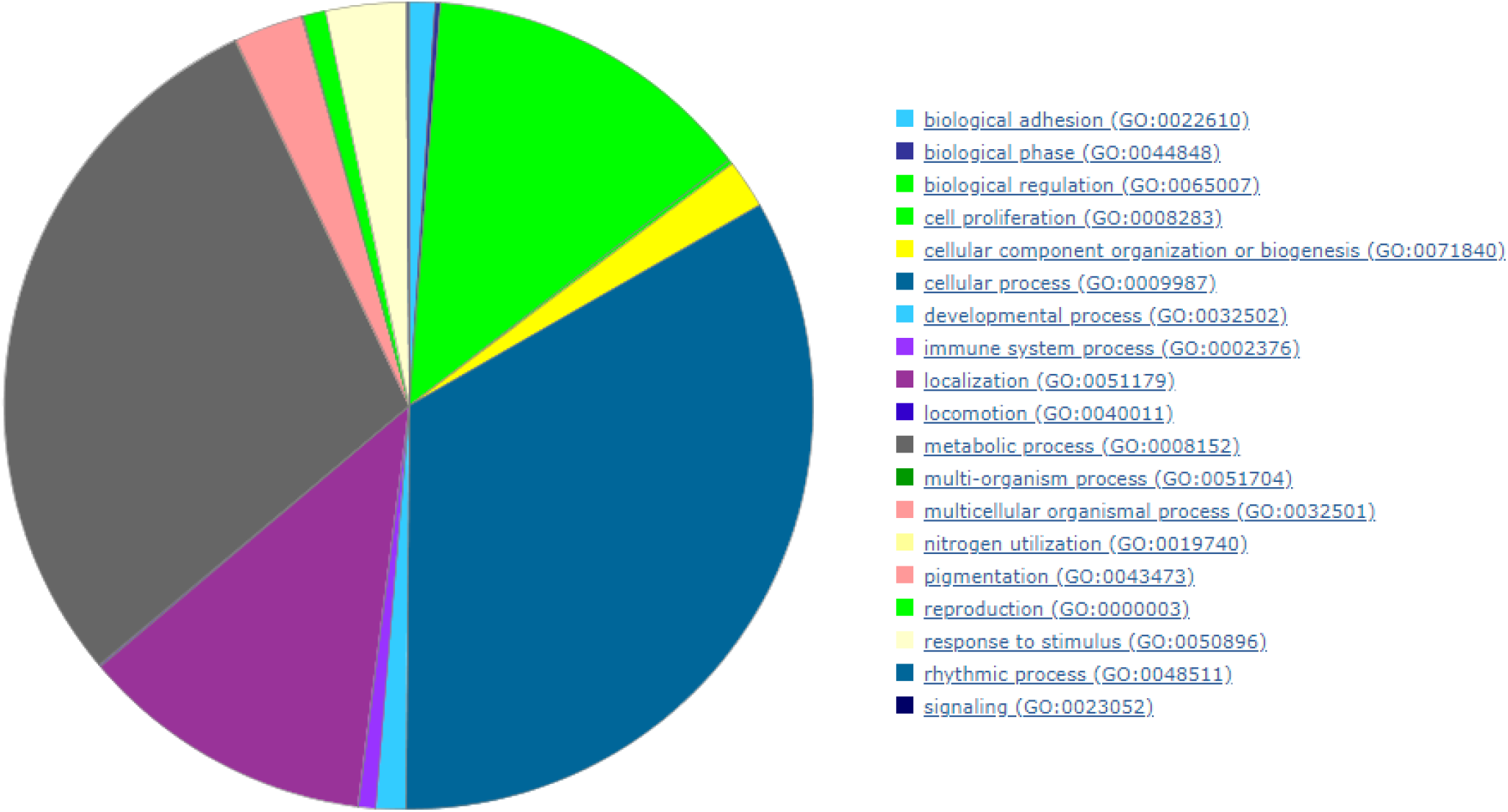
Biological process (Gene Ontology) for PANTHER protein families annotated in *Tubastraea* sp. genome

**Fig. 10.**
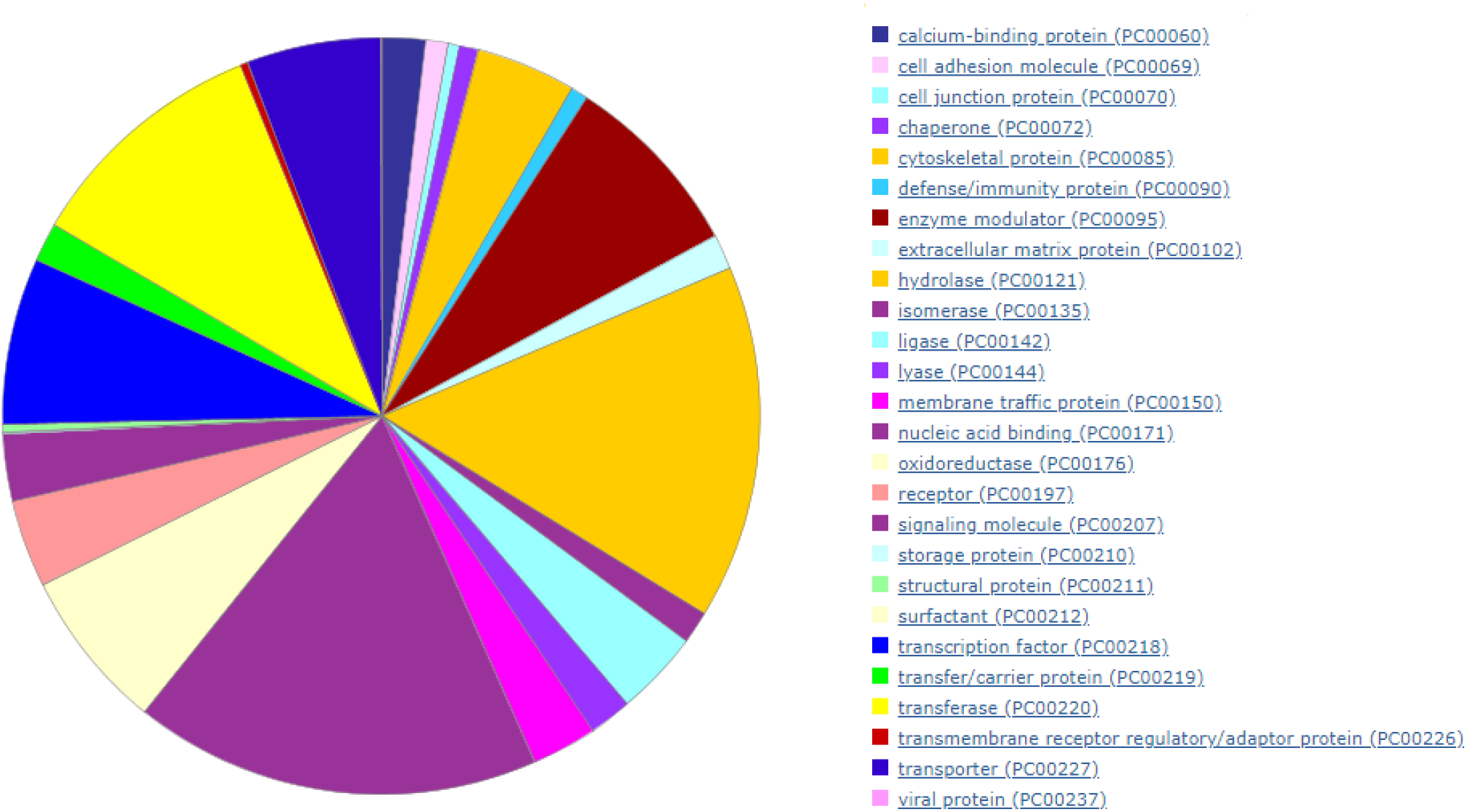
PANTHER protein class for annotated genes in the *Tubastraea* sp. genome

### A.14. Mitochondrial genome assembly and annotation

The assembly using Illumina and PacBio DNA reads generated a single contig containing the whole mtgenome which was compared with the *Tubastraea tagusensis* (NC_030352.1, [50]) sequence. The run with NOVOplasty using RNA Illumina reads also allowed the recovery of the entire mtgenome, except for a single gap region that is located in the region related to the COX1 (also known as COI) intron. The mtgenome annotation using GeSeq found 17 genes, of which 13 encode for proteins and 4 are non-coding genes (Figure 3). When compared with *Tubastraea tagusensis* (NC_030352.1, [50], it has one fewer base due toan indel at position 9792, which is located between the rns and COX3 genes. We also mapped four polymorphic sites that might cause non-synonymous substitution in three proteincoding genes (ND1, *CYTB* and *COX1*) (Table 6).

**Table 6.**
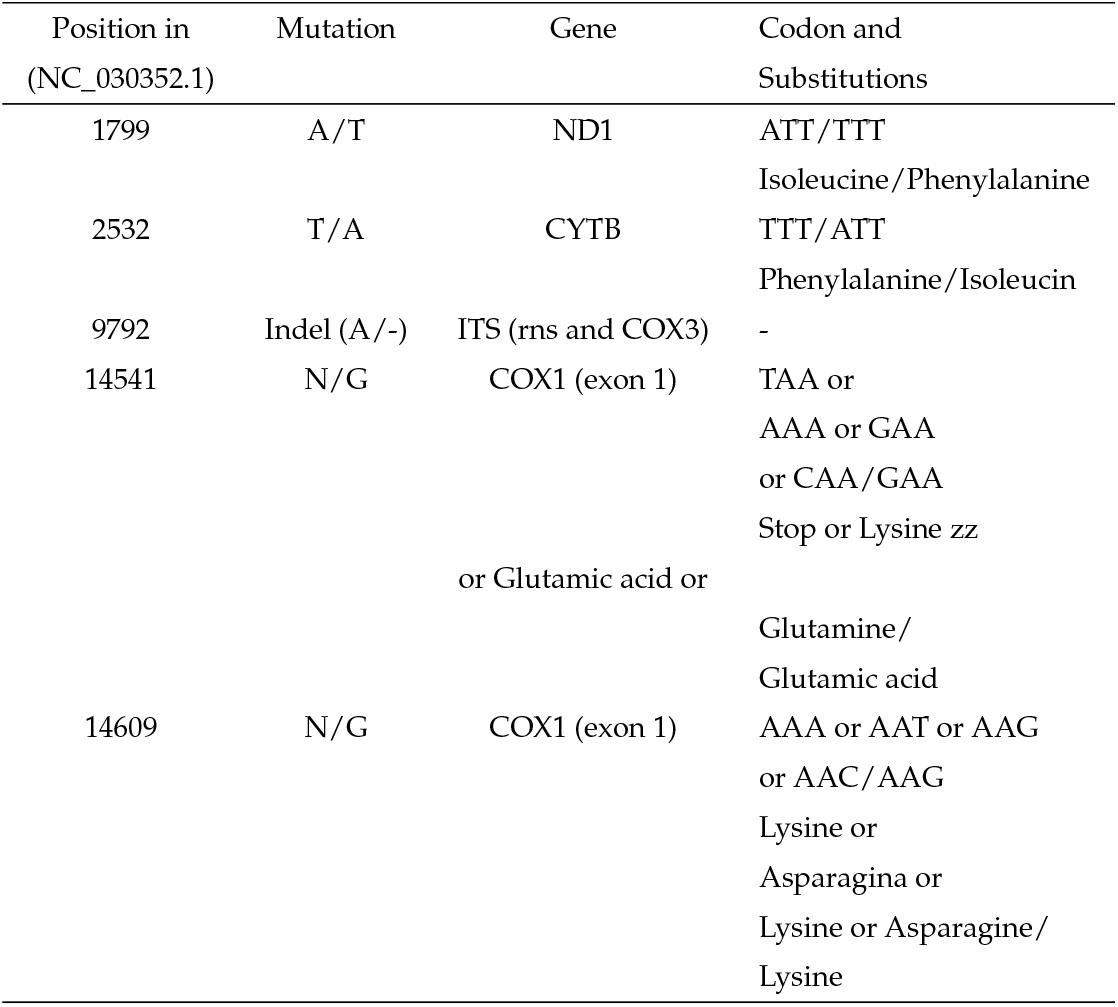
Differences found between NC_030352.1 and the Mtgenome of *Tubastraea* sp. sampled in Angra dos Reis, Brazil through sequence comparison.

| Position in (NC_030352.1) | Mutation | Gene | Codon and Substitutions |
| --- | --- | --- | --- |
| 1799 | A/T | ND1 | ATT/TTT<br>Isoleucine/Phenylalanine |
| 2532 | T/A | CYTB | TTT/ATT<br>Phenylalanine/Isoleucine |
| 9792 | Indel (A/-) | ITS (rns and COX3) | - |
| 14541 | N/G | COX1 (exon 1) | TAA or<br>AAA or GAA<br>or CAA/GAA<br>Stop or Lysine zz<br>or Glutamic acid or<br>Glutamine/<br>Glutamic acid |
| 14609 | N/G | COX1 (exon 1) | AAA or AAT or AAG<br>or AAC/AAG<br>Lysine or<br>Asparagina or<br>Lysine or Asparagine/<br>Lysine |

## REFERENCES

1. Richmond RH, Hunter CL (1990) Reproduction and recruitment of corals: comparisons among the Caribbean, the Tropical Pacific, and the Red Sea. Marine ecology progress series Oldendorf 60(1):185–203.

2. Brusca RC, Brusca GJ (2007) Invertebrados (Guanabara Koogan)

3. White M, Mohn C, de Stigter H, Mottram G (2005) Deep-water coral development as a function of hydrodynamics and surface productivity around the submarine banks of the Rockall Trough, NE Atlantic. Cold-Water Corals and Ecosystems, eds Freiwald A, Roberts JM (Springer Berlin Heidelberg, Berlin, Heidelberg), pp 503–514.

4. Hemond EM, Kaluziak ST, Vollmer SV (2014) The genetics of colony form and function in *Caribbean Acropora* corals. BMC Genomics 15:1133.

5. Luz BLP, et al. (2018) A polyp from nothing: The extreme regeneration capacity of the Atlantic invasive sun corals *Tubastraea coccinea* and *T. tagusensis* (Anthozoa, Scleractinia). J Exp Mar Bio Ecol 503:60–65.

6. Fedders H, Augustin R, Bosch TCG (2004) A Dickkopf-3-related gene is expressed in differentiating nematocytes in the basal metazoan Hydra. Dev Genes Evol 214(2):72–80.

7. Petersen HO, et al. (2015) A Comprehensive Transcriptomic and Proteomic Analysis of Hydra Head Regeneration. Mol Biol Evol 32(8):1928–1947.

8. Bhambri A, et al. (2018) Large scale changes in the transcriptome of *Eisenia fetida* during regeneration. PLoS One 13(9):e0204234.

9. Zhu W, et al. (2012) Retrotransposon long interspersed nucleotide element-1 (LINE-1) is activated during salamander limb regeneration. Dev Growth Differ 54(7):673–685.

10. Lin M-F, et al. (2017) Analyses of Corallimorpharian Transcriptomes Provide New Perspectives on the Evolution of Calcification in the Scleractinia (Corals). Genome Biol Evol 9(1):150–160.

11. Wirshing HH, Baker AC (2014) Molecular evolution of calcification genes in morphologically similar but phylogenetically unrelated scleractinian corals. Mol Phylogenet Evol 77:281–295.

12. Capel KCC, et al. (2017) Clone wars: asexual reproduction dominates in the invasive range of *Tubastraea* spp. (Anthozoa: Scleractinia) in the South-Atlantic Ocean. PeerJ 5:e3873.

13. Shikina S, Chiu Y-L, Lee Y-H, Chang C-F (2015) From Somatic Cells to Oocytes: A Novel Yolk Protein Produced by Ovarian Somatic Cells in a Stony Coral, *Euphyllia ancora*. Biol Reprod 93(3):57.

14. Shikina S, et al. (2013) Yolk formation in a stony coral *Euphyllia ancora* (Cnidaria, Anthozoa): insight into the evolution of vitellogenesis in nonbilaterian animals. Endocrinology 154(9):3447–3459.

15. Vermeij MJA (2005) A novel growth strategy allows *Tubastrea coccinea* to escape small-scale adverse conditions and start over again. Coral Reefs 24(3):442–442.

16. Creed JC, De Paula AF (2007) Substratum preference during recruitment of two invasive alien corals onto shallow-subtidal tropical rocky shores. Mar Ecol Prog Ser 330:101–111.

17. Glynn PW, et al. (2008) Reproductive ecology of the azooxanthellate coral *Tubastraea coccinea* in the Equatorial Eastern Pacific: Part V. Dendrophylliidae. Mar Biol 153(4):529–544.

18. de Paula AF, de Oliveira Pires D, Creed JC (2014) Reproductive strategies of two invasive sun corals (Tubastraea spp.) in the southwestern Atlantic. 94(03)):481–492.

19. Vauguan, W T, Wells, J W (1943) Revision of the suborders, families and genera of the Scleractinia. GSA SPECIAL PAPERS (Geological Society of America), pp 1–363.

20. Creed JC, et al. (2017) The invasion of the azooxanthellate coral *Tubastraea* (Scleractinia: Dendrophylliidae) throughout the world: history, pathways and vectors.

21. Cairns SD (2001) A brief history of taxonomic research on azooxanthellate Scleractinia (Cnidaria: Anthozoa). 7492.

22. Costa TJF, et al. (2014) Expansion of an invasive coral species over Abrolhos Bank, Southwestern Atlantic. Mar Pollut Bull 85(1):252–253.

23. de Oliveira Soares M, Davis M, Carneiro P (2016) Northward range expansion of the invasive coral (*Tubastraea tagusensis*) in the southwestern Atlantic. doi:10.1007/s12526-016-0623-x.

24. de Paula AF, Creed JC (2004) Two species of the coral Tubastraea (Cnidaria, Scleractinia) in Brazil: A case of accidental introduction. 74(1)):175–183.

25. Ferreira CEL (2003) Non-indigenous corals at marginal sites. 22(4)):498–498.

26. Mantelatto MC, Creed JC, Mourão GG, Migotto AE, Lindner A (2011) Range expansion of the invasive corals *Tubastraea coccinea* and *Tubastraea tagusensis* in the Southwest Atlantic. Coral Reefs 30(2):397–397.

27. Sampaio CLS, Miranda RJ, Maia-Nogueira R, Nunes J de ACC (2012) New occurrences of the nonindigenous orange cup corals *Tubastraea coccinea* and *T. tagusensis* (Scleractinia: Dendrophylliidae) in Southwestern Atlantic. CheckList 8(3):528–530.

28. Silva AG, et al. (2011) Expansion of the invasive corals *Tubastraea coccinea* and *Tubastraea tagusensis* into the Tamoios Ecological Station Marine Protected Area, Brazil. Aquat Invasions 6(Suppl 1):S105–S110.

29. Brito A, López C, Vicente OO, Herrera R, Cuervo JJS (2017) Colonización y expansión en Canarias de dos corales potencialmente invasores introducidos por las plataformas petrolíferas. 45:65–82.

30. López C, et al. (2019) Invasive *Tubastraea* spp. and *Oculina patagonica* and other introduced scleractinians corals in the Santa Cruz de Tenerife (Canary Islands) harbor: Ecology and potential risks. Regional Studies in Marine Science 29:100713.

31. Ocaña O, et al. (2015) A survey on Anthozoa and its habitats along the Northwest African coast and some islands: new records, descriptions of new taxa and biogeographical, ecological and taxonomical comments. Part I. Available at: http://rua.ua.es/dspace/handle/10045/52860.

32. Shinzato C, et al. (2011) Using the *Acropora digitifera* genome to understand coral responses to environmental change. Nature 476(7360):320–323.

33. Ying H, et al. (2019) The Whole-Genome Sequence of the Coral *Acropora millepora*. Genome Biol Evol 11(5):1374–1379.

34. Helmkampf M, Bellinger MR, Geib SM, Sim SB, Takabayashi M (2019) Draft Genome of the Rice Coral *Montipora capitata* Obtained from Linked-Read Sequencing. Genome Biol Evol 11(7):2045–2054.

35. Prada C, et al. (2016) Empty Niches after Extinctions Increase Population Sizes of Modern Corals. Curr Biol 26(23):3190–3194.

36. Cunning R, Bay RA, Gillette P, Baker AC, Traylor-Knowles N (2018) Comparative analysis of the *Pocillopora damicornis* genome highlights role of immune system in coral evolution. Sci Rep 8(1):16134.

37. Celis JS, et al. (2018) Binning enables efficient host genome reconstruction in cnidarian holobionts. Gigascience 7(7). doi:10.1093/gigascience/giy075.

38. Voolstra CR, et al. (2017) Comparative analysis of the genomes of *Stylophora pistillata* and *Acropora digitifera* provides evidence for extensive differences between species of corals. Sci Rep 7(1):17583.

39. Wells JW (1982) Notes on Indo-Pacific Scleractinian Corals. Part 9. New Corals from the Galapagos Islands. Available at: https://scholarspace.manoa.hawaii.edu/handle/10125/421 [Accessed August 28, 2019].

40. Cairns SD, Zibrowius H (1997) Cnidaria Anthozoa: azooxanthellate Scleractinia from the Philippine and Indonesian regions. 7876.

41. Dolezel J, Greilhuber J, Suda J (2007) Estimation of nuclear DNA content in plants using flow cytometry. Nat Protoc 2(9):2233–2244.

42. Garcia GD, et al. (2013) Metagenomic analysis of healthy and white plague-affected *Mussismilia braziliensis* corals. Microb Ecol 65(4):1076–1086.

43. Anderson DA, Walz ME, Weil E, Tonellato P, Smith MC (2016) RNA-Seq of the Caribbean reef-building coral *Orbicella faveolata* (Scleractinia-Merulinidae) under bleaching and disease stress expands models of coral innate immunity. PeerJ 4:e1616.

44. Wingett SW, Andrews S (2018) FastQ Screen: A tool for multi-genome mapping and quality control. F1000Res 7:1338.

45. Mapleson D, Accinelli GG, Kettleborough G, Wright J, Clavijo BJ (2016) KAT: a K-mer analysis toolkit to quality control NGS datasets and genome assemblies. Bioinformatics:btw663.

46. Zimin AV, et al. (2013) The MaSuRCA genome assembler. Bioinformatics 29(21):2669–2677.

47. Gurevich A, Saveliev V, Vyahhi N, Tesler G (2013) QUAST: quality assessment tool for genome assemblies. Bioinformatics 29(8):1072–1075.

48. Waterhouse RM, et al. (2017) BUSCO applications from quality assessments to gene prediction and phylogenomics. Mol Biol Evol. doi:10.1093/molbev/msx319.

49. Roach MJ, Schmidt SA, Borneman AR (2018) Purge Haplotigs: allelic contig reassignment for third-gen diploid genome assemblies. BMC Bioinformatics 19(1):460.

50. Capel KCC, et al. (2016) Complete mitochondrial genome sequences of Atlantic representatives of the invasive Pacific coral species *Tubastraea coccinea* and *T. tagusensis* (Scleractinia, Dendrophylliidae): Implications for species identification. Gene 590(2):270–277.

